# Non-neural factors influencing BOLD response magnitudes within individual subjects

**DOI:** 10.1101/2021.12.26.474185

**Authors:** Jan W. Kurzawski, Omer Faruk Gulban, Keith Jamison, Jonathan Winawer, Kendrick Kay

## Abstract

To what extent is the size of the blood-oxygen-level-dependent (BOLD) response influenced by factors other than neural activity? In a re-analysis of three neuroimaging datasets (male and female human participants), we find large systematic inhomogeneities in the BOLD response magnitude in primary visual cortex (V1): stimulus-evoked BOLD responses, expressed in units of percent signal change, are up to 50% larger along the representation of the horizontal meridian than the vertical meridian. To assess whether this surprising effect can be interpreted as differences in local neural activity, we quantified several factors that potentially contribute to the size of the BOLD response. We find relationships between BOLD response magnitude and cortical thickness, curvature, depth and macrovasculature. These relationships are consistently found across subjects and datasets and suggest that variation in BOLD response magnitudes across cortical locations reflects, in part, differences in anatomy and vascularization. To compensate for these factors, we implement a regression-based correction method and show that after correction, BOLD responses become more homogeneous across V1. The correction reduces the horizontal/vertical difference by about half, indicating that some of the difference is likely not due to neural activity differences. We conclude that interpretation of variation in BOLD response magnitude across cortical locations should consider the influence of the potential confounding factors of thickness, curvature, depth and vascularization.

**Significance statement:** The magnitude of the BOLD signal is often used as a surrogate of neural activity, but the exact factors that contribute to its strength have not been studied on a voxel-wise level. Here, we examined several anatomical and measurement-related factors to assess their relationship with BOLD signal magnitude. We find that BOLD magnitude correlates with cortical anatomy, depth and macrovasculature. To remove the contribution of these factors, we propose a simple, data-driven correction method that can be used in any functional magnetic resonance imaging (fMRI) experiment. After accounting for the confounding factors, BOLD magnitude becomes more spatially homogenous. Our correction method improves the ability to make more accurate inferences about local neural activity from fMRI data.

## Introduction

The blood-oxygen-level-dependent (BOLD) signal measured by fMRI is an important tool for non-invasive study of the human nervous system. However, the neural mechanisms underlying BOLD remain an active area of investigation (Herman et al., 2017). One clear conclusion is that the BOLD signal is strongly influenced by neural activity (Arthurs et al., 2000; Heeger et al., 2000; Attwell and Iadecola, 2002; Heeger and Ress, 2002; Logothetis, 2002; Lee et al., 2010; Siero et al., 2014). For a given location in the brain, and within a constrained paradigm (e.g., viewing different images and measuring the response that they elicit in visual cortex), the BOLD signal magnitude appears to be lawfully related to basic measures of neural activity. For example, as stimulus contrast increases, neural firing rates and BOLD magnitude increase in proportion (Heeger et al., 2000). Similarly, increase in coherence of stimulus motion boosts BOLD magnitude and firing rates in V5/MT (Britten et al., 1993; Rees et al., 2000). When comparing different experimental paradigms or different brain locations, however, it is less clear how to interpret differences in the magnitude of the BOLD signal. For example, seeing a stimulus and expecting a stimulus can both elicit robust BOLD signals in V1, but the underlying neural activity is very different in the two paradigms (Sirotin and Das, 2009; Herman et al., 2017). It is also the case that similar BOLD signal magnitudes in two locations may be linked to very different underlying neural activity. These two limitations are reviewed by (Logothetis, 2008).

There are several reasons to believe that BOLD signal magnitudes, even within a fixed experimental paradigm, are influenced by factors that are not directly related to neural activity. The BOLD response, quantified in terms of percent signal change, can be especially high in voxels containing large veins (Menon et al., 1993; Kim et al., 1994; Hoogenraad et al., 1999; Kay et al., 2019) or unusually low, delayed, and/or displaced in voxels near cerebral sinuses (Winawer et al., 2010; Jamison et al., 2017). The choice of MRI sequence, field strength (van der Zwaag et al., 2009), and sequence parameters like echo time (Gorno-Tempini et al., 2002) can also affect BOLD signal magnitude, and these effects may vary across the brain (Herman et al., 2017). Indeed, it has been reported that BOLD may vary across the cortex up to 40% simply due to different orientation of vasculature relative to the direction of the static magnetic field (Gagnon et al., 2015a; Gagnon et al., 2016; Viessmann et al., 2019). Furthermore, recent high-resolution fMRI studies have shown that BOLD signal magnitude clearly depends on cortical depth. It is highest in the superficial depths which are positioned near large pial veins and decreases with depth (Polimeni et al., 2010; Koopmans et al., 2011; Zimmermann et al., 2011; Yu et al., 2014; Fracasso et al., 2016a; Fracasso et al., 2016b; Dumoulin, 2017; Dumoulin et al., 2018; Kay et al., 2019; Self et al., 2019; van Dijk et al., 2020).

In this paper, we study variations in BOLD signal magnitude within a fixed paradigm, focusing our efforts on primary visual cortex (V1). We believe that by focusing on a single brain region in well-controlled visual paradigms, we are in the best position to derive sound interpretations of differences in BOLD signal magnitudes across the cortex. In three distinct datasets, we demonstrate large differences between the meridian locations: the BOLD magnitude in V1 is up to 50% higher along the representation of the horizontal meridian than along the representation of the vertical meridian. We then investigate the potential basis of these inhomogeneities by analyzing factors that are in principle distinct from neural activity. As non-neural factors we consider cortical curvature, cortical thickness, cortical depth, presence of macrovasculature (as indexed by bias-corrected EPI intensity), angle with respect to *B*_0_ magnetic field and radiofrequency (RF) coil bias. We motivate the selection of these factors in the Methods. We find that several of these factors are systematically related to observed variation in BOLD magnitudes across V1. To remove their influence, we propose a simple correction method and show that the correction increases BOLD signal homogeneity across V1, reducing the difference in response across the horizontal and vertical meridians by about half.

## Methods

### Datasets

We used three publicly available visual fMRI datasets: the Human Connectome Project (HCP) 7T Retinotopy Dataset (Benson et al., 2018), the Natural Scenes Dataset (NSD) (Allen et al., 2021), and the Temporal Decomposition Method (TDM) Dataset (Kay et al., 2020). All data were acquired on 7T MR scanners using gradient-echo pulse sequences (technical details provided in Table 1). The datasets varied in stimulus properties and experimental design. HCP stimuli consisted of rings, wedges, and bars in a retinotopic mapping experiment; NSD stimuli consisted of natural scene images; and TDM stimuli consisted of high-contrast rings presented at different eccentricities. Experimental details are shown in **Figure 1**. The analyses performed in this paper start with pre-processed data from each dataset.

**Table 1.** Details on the fMRI pulse sequence used in each of the datasets. Each column describes different dataset.

| Dataset | TDM | NSD | HCP |
| --- | --- | --- | --- |
| Field strength | 7T | 7T | 7T |
| TR | 2200 ms | 1600 ms | 1000 ms |
| TE | 22.4 ms | 22.0 ms | 22.2 ms |
| Flip angle | 80 | 62 | 45 |
| Number of slices | 84 | 84 | 85 |
| Matrix size | $200 \times 162$ | $120 \times 120$ | $130 \times 130$ |
| Field of view | $160 \text{ mm} \times 129.6 \text{ mm}$ | $216 \text{ mm} \times 216 \text{ mm}$ | $208 \text{ mm} \times 208 \text{ mm}$ |
| **Nominal spatial resolution** | 0.8 mm | 1.8 mm | 1.6 mm |
| **Multiband factor** | 2 | 3 | 5 |
| **iPAT factor** | 3 | 2 | 2 |
| **Partial Fourier** | 6/8 | 7/8 | 7/8 |

**Figure 1.**
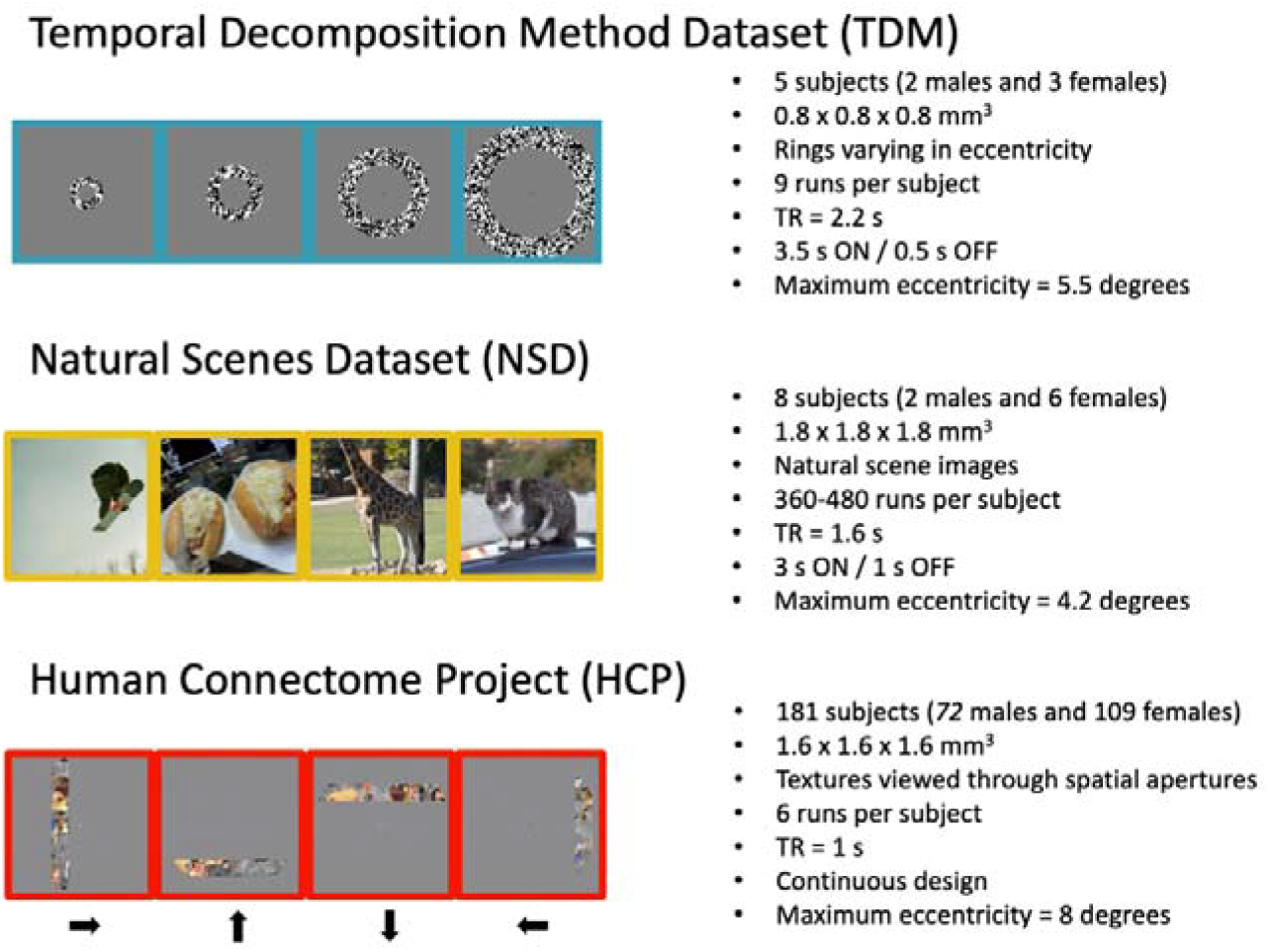
Datasets used in this study. Stimulus images for each of the datasets are shown. For TDM, stimuli consisted of 6 rings varying in eccentricity. For NSD, stimuli consisted of natural scene images. For HCP, the experiment consisted of several retinotopic mapping runs that included expanding and contracting rings, rotating wedges, and moving bars filled with a colorful object-based texture. Additional acquisition details are provided in Table 1.

### Extracting BOLD magnitude

From each dataset, we extracted a measure of BOLD signal magnitude at each cortical surface vertex. For TDM, we started with the pre-processed fMRI time-series data provided with the dataset and analyzed the data with a GLM. Specifically, we convolved a canonical HRF with stimulus onsets to create a regressor for each experimental condition, and then used these regressors with GLMdenoise (Kay et al., 2013b) to estimate a beta weight for each condition. We computed the maximum beta weight across all conditions for each voxel as the measure of BOLD signal magnitude. These results are defined at six different depths (equidistant from 10% to 90% of the cortical thickness) in each subject’s native surface space. (Depth assignment was achieved by a spatial interpolation of each fMRI volume at the locations of the six depth-dependent cortical surfaces; see Kay et al. (2020) for details.) For NSD, we took the ‘meanbeta’ values (1 mm data preparation, beta version 2) provided with the dataset; these values indicate the average BOLD percent signal change observed across all stimulus trials and all scan sessions. We then mapped these values to the 3 depth surfaces provided in NSD (positioned at 25%, 50%, and 75% of the cortical thickness). The HCP dataset was previously analyzed (Benson et al., 2018) with a population receptive field (pRF) model (Dumoulin and Wandell, 2008) implemented in analyzePRF (Kay et al., 2013a). The model includes a gain parameter that describes the amplitude of the BOLD response of a given voxel (or vertex) to the object-based texture (covering the entire pRF) for a single repetition time (TR = 1 s). We quantified BOLD in terms of percent signal change (%BOLD) by dividing the gain parameter by mean signal intensity and multiplying by 100. The results are prepared in FreeSurfer’s fsaverage space.

### Visual field mapping

We used retinotopic mapping to divide the primary visual cortex into a set of regions. For HCP, we used polar angle and eccentricity estimates available from the data release. For the TDM and NSD datasets, we mapped Benson’s polar angle and eccentricity atlas using neuropythy software (Benson and Winawer, 2018). We use the following convention for all 3 datasets: the upper vertical meridian corresponds to 0 deg, the horizontal meridian corresponds to 90 deg, and the lower vertical meridian corresponds to 180 deg. Note that the polar angle estimates are rescaled for the correlation and linear regression analysis (see next section). We used Benson’s definition of the extent of visual areas V1, V2, and V3 for all 3 datasets (Benson et al., 2014).

### Quantification of non-neural factors

In the TDM and NSD datasets, we quantified several factors that might be related to variation in the magnitude of the BOLD signal across cortical locations. We focused on factors that can be easily extracted from either functional or anatomical data that are typically acquired in an fMRI experiment. For the purposes of the present study, we consider only within-subject factors rather than across-subject factors, with the goal of removing non-neural influences on the variation of BOLD magnitudes across voxels. We note that there are several other factors that influence variation of overall BOLD magnitude across subjects like caffeine use (Liu et al., 2004), vascular age (Tsvetanov et al., 2021), and heart rate (Chang et al., 2009). Below, we describe each of the within-subject factors that we considered in the present study.

*Curvature* was obtained from FreeSurfer outputs (Dale et al., 1999; Fischl and Dale, 2000), and refers to the geometry of the folding pattern of the cortical surface. Negative values correspond to gyri while positive values correspond to sulci. Curvature is quantified as 1/r, where *r* is the radius of an inscribed circle measured in mm.

*Thickness* was also obtained from FreeSurfer outputs. It is measured in mm and corresponds to the distance between the outermost (close to cerebrospinal fluid) and innermost (close to white matter) boundaries of gray matter. Curvature and thickness are well known to vary across visual cortex. Their relationship with %BOLD remains unknown and has not been investigated in detail, especially on a voxel-by-voxel basis. We include these factors in our analysis to assess whether these anatomical factors have systematic relationships with BOLD magnitude.

*Mean bias-corrected EPI* was calculated as the mean signal intensity in the fMRI data divided by the estimated RF coil bias (details below). The units range from approximately 0 to 2, and indicate percentages (e.g., 0.5 means 50% of the strength of typical signal intensities). Mean bias-corrected EPI values can be viewed as high spatial frequency changes in signal intensity across space. We include this factor in the analysis as mean bias-corrected EPI was previously found to be a good predictor for venous effects (Kay et al., 2019). Proximity to veins often results in increased BOLD magnitude.

*Depth* was estimated by generating 6 cortical surfaces (for TDM) or 3 cortical surfaces (for NSD) equally spaced between 10% and 90% (for TDM) or 25% and 75% (for NSD) of the distance from the pial surface to the boundary between gray and white matter. These surfaces are numbered from 1 to *n*, where 1 is outermost and *n* is innermost. We include depth as a factor as it is well known that BOLD magnitude is highest in superficial depths and decreases towards the white matter (Polimeni et al., 2010).

*Angle with respect to B*_0_ was calculated by considering the angle (*theta*) between the pial surface normal and the direction of the *B*_0_ static magnetic field as estimated from NIFTI header information. Angle was quantified in degrees and was normalized as abs(*theta*–90) such that a final value of 0 deg indicates that the cortical surface is parallel to the magnetic field and a final value of 90 deg indicates that the cortical surface is perpendicular to the magnetic field. We include angle with respect to *B*_0_ in the analysis because previous reports showed that the BOLD magnitude varies with *B*_0_ angle (Gagnon et al., 2015b).

*RF coil bias* was taken to be the result of fitting a 3D polynomial to the mean signal intensity in the fMRI data. The values are in raw scanner units and represent low spatial frequency changes in the intensity of voxels. This estimation method has been used previously (Kay et al., 2019). We include RF coil bias as a control in our analysis. In theory, there should not be a systematic relationship between RF coil bias and BOLD magnitude, as we express BOLD magnitudes at each voxel in terms of percent signal change (as is typically done in the field), and percent signal change is sensitive to an overall scale factor on the signal.

In sum, all of these factors are known to vary across the cortical surface of V1. The exact biophysical mechanisms that might explain their impact on %BOLD are in some cases unknown (e.g., curvature). In other cases, we expect that some factors should not bear systematic relationships to %BOLD (e.g., RF coil bias). In general, the work here is intended to be a first step towards understanding the influence of potential non-neural contributions to variations in %BOLD across individual voxels within a given subject.

### Quantification of neural factors

*Polar angle* was obtained from Benson’s atlas (Benson et al., 2014), representing the visual field angle to which each cortical location is optimally tuned. For the purposes of our analyses, we normalize polar angle such that 0 deg corresponds to the horizontal meridian and 90 deg corresponds to the upper and lower vertical meridians. We include polar angle as a positive control: we expect that polar angle should bear a systematic relationship with BOLD magnitude, as this is the original observation that motivated the present study.

### Definition of regions of interest

Using the visual field mapping results, we defined regions of interest (ROIs) corresponding to the representation of the horizontal and vertical meridians within V1. The ROIs were defined by limiting the eccentricity to the maximum stimulus eccentricity used in each dataset and limiting the angle to a specific range (e.g., to create a V1 ROI for the upper vertical meridian with a width of 20 deg, we created a mask where polar angle estimates were higher than 0 and lower than 20).

### Modelling variation in BOLD signal magnitude

To account for non-neural contribution to %BOLD, we used a multiple regression model. The modeled data (Y) consisted of the %BOLD value observed at each surface vertex in visual areas V1–V3. Although this study focuses on BOLD homogeneity in V1, we include %BOLD in V1–V3. This is because we are attempting to establish relationships that might generalize across different cortical regions. Furthermore, if we were to include only vertices in V1, we would be at high risk of removing genuine neural activity differences (e.g. those that may exist between the horizontal and vertical meridians) that correlate with the non-neural factors.

The variables used to model the data included thickness, curvature, depth and mean bias-corrected EPI intensity. (Only these four factors showed evidence of being substantially related to BOLD magnitude; see Results.) The variables were standardized (z-scored) and, together with a constant term, were included as predictors in the design matrix (X). Ordinary least-squares estimates for beta weights were obtained in the following linear model:

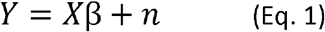

where *Y* is the %BOLD magnitude at each vertex, *X* is the 5-column design matrix, *β* is a set of beta weights (5 per vertex), and *n* is a set of residuals.

### Major cortical sulci

In several figures we show outlines of major cortical sulci. These include the calcarine sulcus (CALC), parieto-occipital sulcus (POS), intraparietal sulcus (IPS), occipitotemporal sulcus (OTS), and superior temporal sulcus (STS). These sulci were manually labelled on the fsaverage surface and then mapped to each individual’s native surface.

### Data and code availability

The datasets used in this paper are freely available online: NSD (http://naturalscenesdataset.org), HCP (https://osf.io/bw9ec/), and TDM (https://osf.io/j2wsc/). Code that reproduces the main figures in this paper is available at https://github.com/jk619/meridianbias/. Associated data files are available at https://osf.io/2nc4x/.

## Results

### Stronger BOLD responses along the V1 horizontal meridian

We examined BOLD response magnitudes in three freely available datasets: the Natural Scenes Dataset (NSD; Allen et al., 2021), the data used for the Temporal Decomposition Method (TDM; Kay et al., 2020), and the Human Connectome Project 7T Retinotopy Dataset (HCP; Benson et al., 2018). Each dataset contains BOLD responses to different types of visual stimulation (see Methods). We defined one region of interest (ROI) for the horizontal meridian (HM) and one for the vertical meridian (VM) **(Figure 2A–B)**. These ROIs represent a wedge-shaped region in the visual field centered at the horizontal meridian with a width of 40 deg (horizontal) and two wedges abutting the vertical meridian each with a width of 20 deg (vertical).

**Figure 2.**
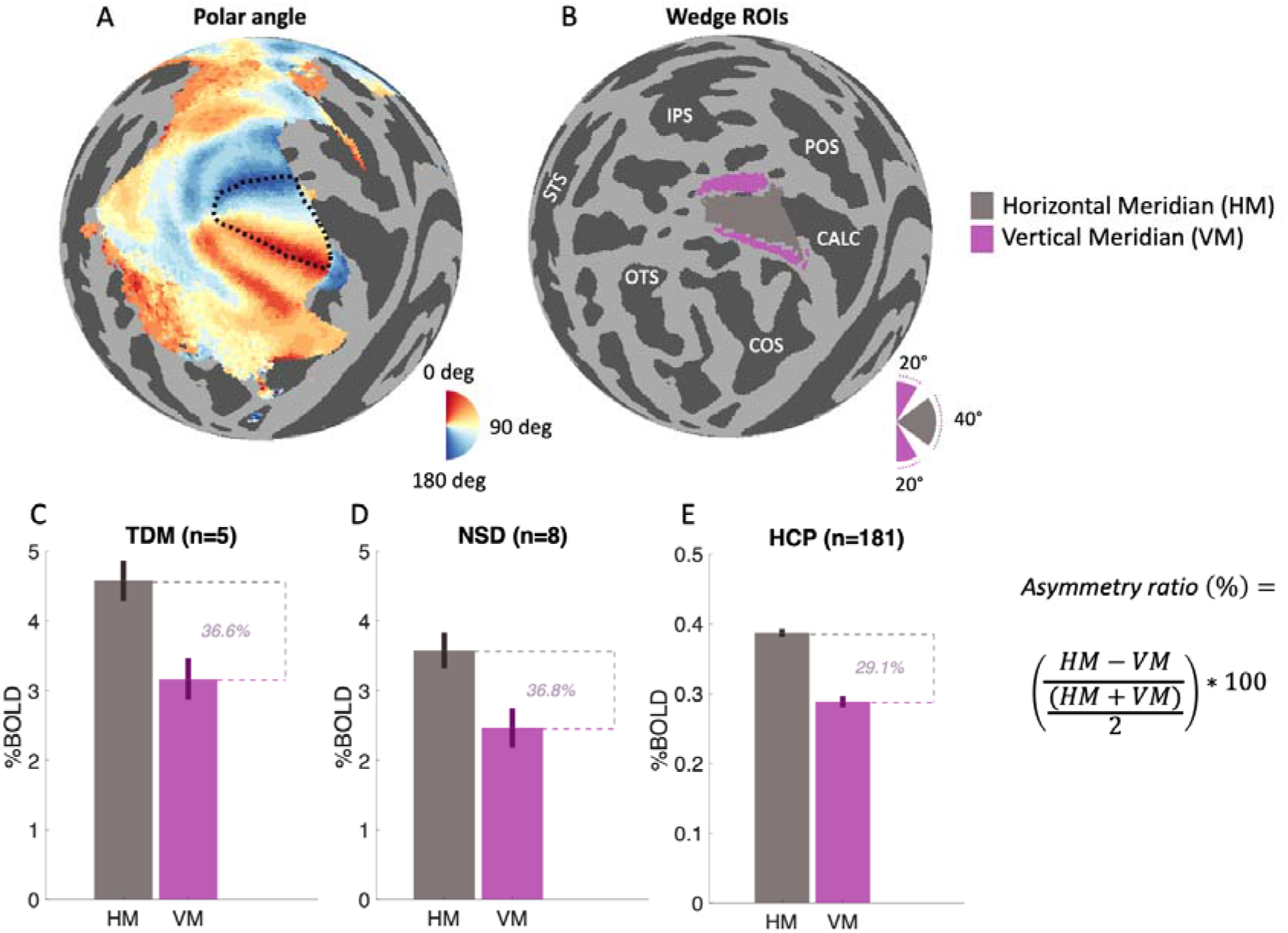
BOLD magnitude is higher at the horizontal meridian in V1. **A)** Polar angle map of group-average HCP subject (999999) with V1 boundary outlined in dotted black lines. **B)** Horizontal and vertical regions of interest (ROIs) are indicated in gray and magenta, respectively. White text indicates major brain sulci (see Methods). **C-E)** Mean BOLD magnitude for horizontal and vertical ROIs in the three datasets. Error bars indicate standard error across subjects.

In each of the three datasets, we compared BOLD magnitudes expressed in percent signal change (%BOLD) observed for the VM with BOLD magnitudes observed for the HM **(Figure 2C–E)**. In each dataset, we find higher %BOLD in the HM ROIs compared to the VM ROIs. We summarize this difference with an asymmetry ratio: (HM–VM)/mean(HM,VM). All datasets show strong asymmetry, with an asymmetry ratio of ~30%. Positive values for the asymmetry ratio indicate greater response for the horizontal meridian. (Note that if the asymmetry is expressed as a percentage of the smaller vertical meridian response, the increase reflected in the larger horizontal meridian response is up to ~50%.)

One possibility is that the horizontal and vertical V1 BOLD responses are in fact similar, but the vertical ROIs appear to have lower signal due to mixing with signal from V2. V2 and V1 border along the vertical meridian representation, and blurring might occur either in acquisition or in pre-processing and analysis. To further our understanding of the V1 response asymmetries, we re-computed asymmetry ratios using smaller wedges at many locations **(Figure 3A)**. Note that, because we use smaller wedges, the asymmetry at the cardinal meridians is different from **Figure 2**. While the asymmetry is strongest at the cardinal meridians, some horizontal/vertical asymmetry is found at least 30 deg away from the meridians in all three datasets **(Figure 3B)**. This argues against the explanation that the asymmetry is caused by spillover from V2.

**Figure 3.**
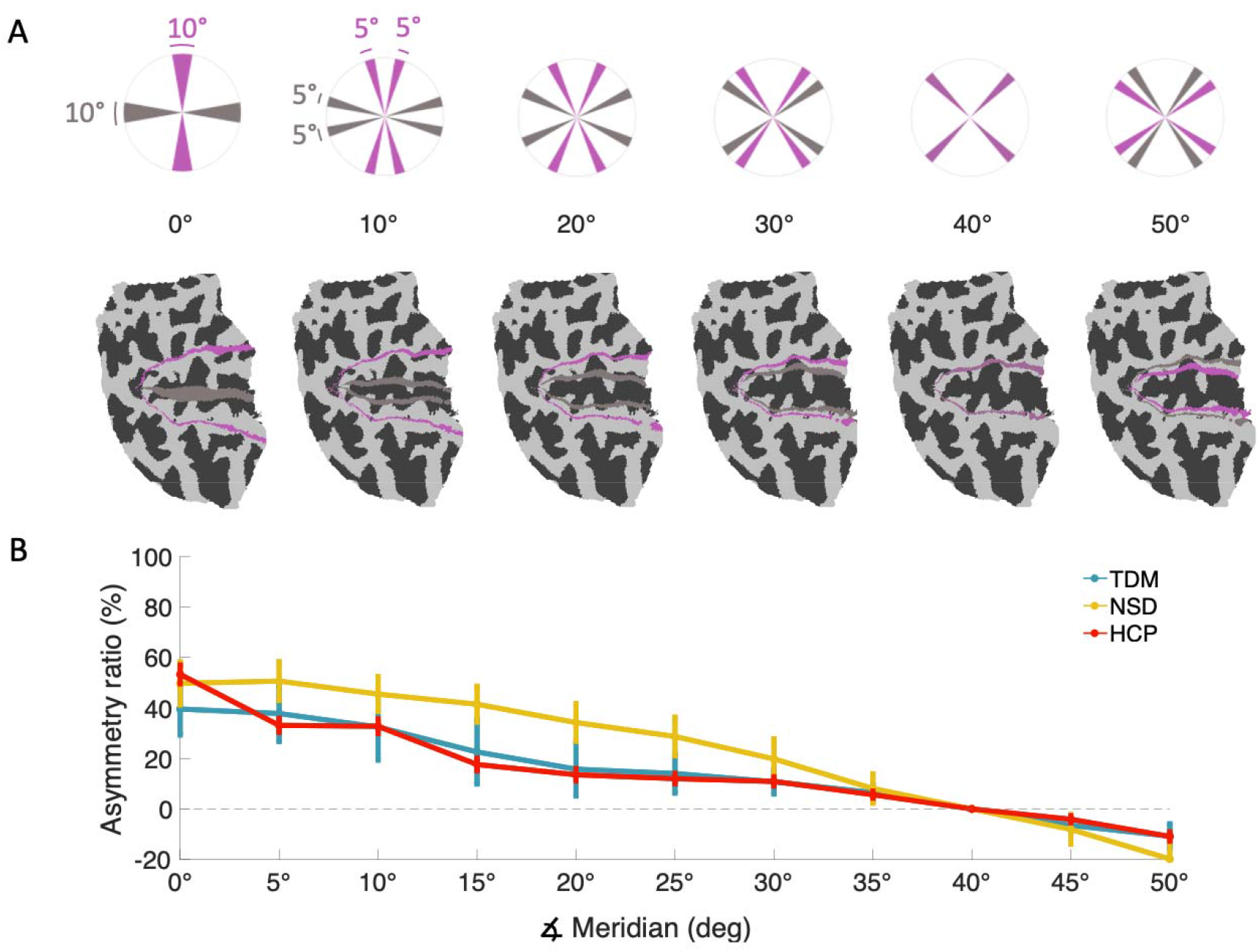
BOLD asymmetries generalize to off-cardinal locations. To further understand V1 BOLD asymmetry, we manipulated the location of the wedge ROIs in 5-deg increments. **A)** The upper row shows the visual field location of wedge ROIs and the lower row shows the corresponding cortical locations (flattened left hemisphere). For clarity, we show only every other set of ROIs. **B)** Asymmetry ratio as a function of angular distance from the cardinal meridians. Error bars indicate standard error across subjects.

### HM/VM asymmetry persists at inner cortical depths

The BOLD signal is strongly influenced by properties of the brain’s vasculature. Uneven venous contributions across the brain can cause variation in BOLD magnitude (Menon et al., 1993; Kim et al., 1994; Hoogenraad et al., 1999; Kay et al., 2019). One possibility is that the meridian asymmetries we observe arise from non-uniformities in the vascular network. To investigate this possibility, we took advantage of the sub-millimeter resolution of the TDM dataset and examined HM/VM asymmetry as a function of depth. Because macroscopic venous effects are larger in the superficial cortex due to large pial veins (Duvernoy et al., 1981; Turner, 2002; Polimeni et al., 2010; Kay et al., 2019), by sampling BOLD responses from deeper depths, we minimize contributions from pial veins. We find that the HM/VM asymmetry is larger at the superficial depths, suggesting that part of the asymmetry may be due to differential properties in macroscopic vasculature **(Figure 4)**. This depth effect is systematic: every subject shows higher asymmetry at the superficial depth than the middle depth. Nonetheless, there remains a substantial horizontal/vertical asymmetry at all depths **(Figure 4)**, suggesting that macroscopic vessels near the pial surface are not the entire explanation. At the innermost depth sampled, which is least influenced by pial vessels, the HM/VM asymmetry is 26% (average across subjects) and is positive in each of the 5 subjects. The middle depths appear to have the least asymmetry. This could be due to a difference in neural responses at intermediate depths, which generally correspond to input-related cortical layers.

**Figure 4.**
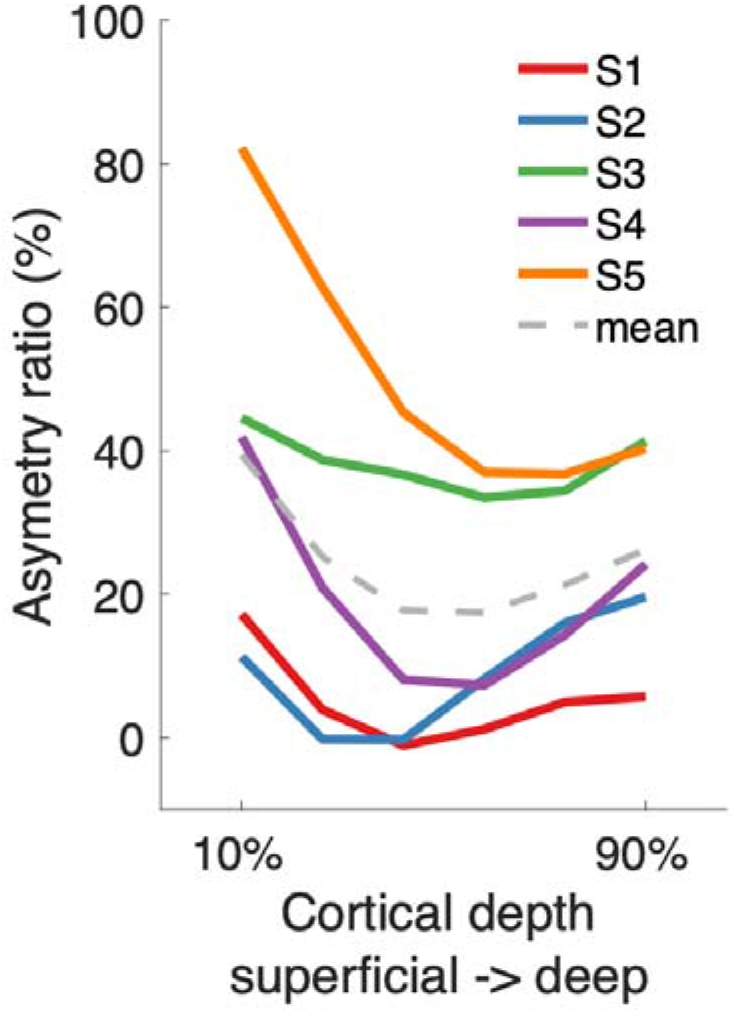
BOLD asymmetry in V1 persists at inner depths. We exploit the high-resolution TDM dataset to discriminate V1 BOLD responses across depth and estimate response asymmetries as a function of depth (asymmetry is calculated in the same way as in **Figure 3)**. The presence of asymmetry at the innermost depth suggests that response asymmetries exist even with minimal contribution of large pial veins.

### Assessing and modeling non-neural contributions to BOLD signal magnitude

In addition to vascular effects, other factors unrelated to neural activity evoked by the experimental manipulation may influence variation in %BOLD across the cortical surface. These additional factors are often neglected in fMRI analysis pipelines. Although some of the factors are known to vary across the cortex, their influence on the BOLD signal is poorly understood. Here, we attempt to understand how these factors may be related to BOLD magnitude variations. To the best of our knowledge, we are unaware of any previous study that has examined this issue in detail, especially at the level of individual voxels (or vertices) within individual subjects.

We first identified a list of possible confounding factors (beyond cortical depth, which we have already introduced) based on consideration of basic anatomical properties of the brain and the nature of fMRI measurement. These factors are cortical curvature, cortical thickness, RF coil bias, mean bias-corrected EPI signal intensity, and angle with respect to *B*_0_. Each of these factors can be interpreted as spatial maps, with a value at each vertex on the cortical surface mesh. The five maps can be obtained from standard anatomical scans (T1-weighted) or from the fMRI measurements themselves without additional MRI experiments (see Methods for details). Example surface visualizations of these maps together with %BOLD are shown in **Figure 5**. We hypothesize that inhomogeneities in some of these maps might explain some of the observed inhomogeneity in %BOLD across V1.

**Figure 5.**
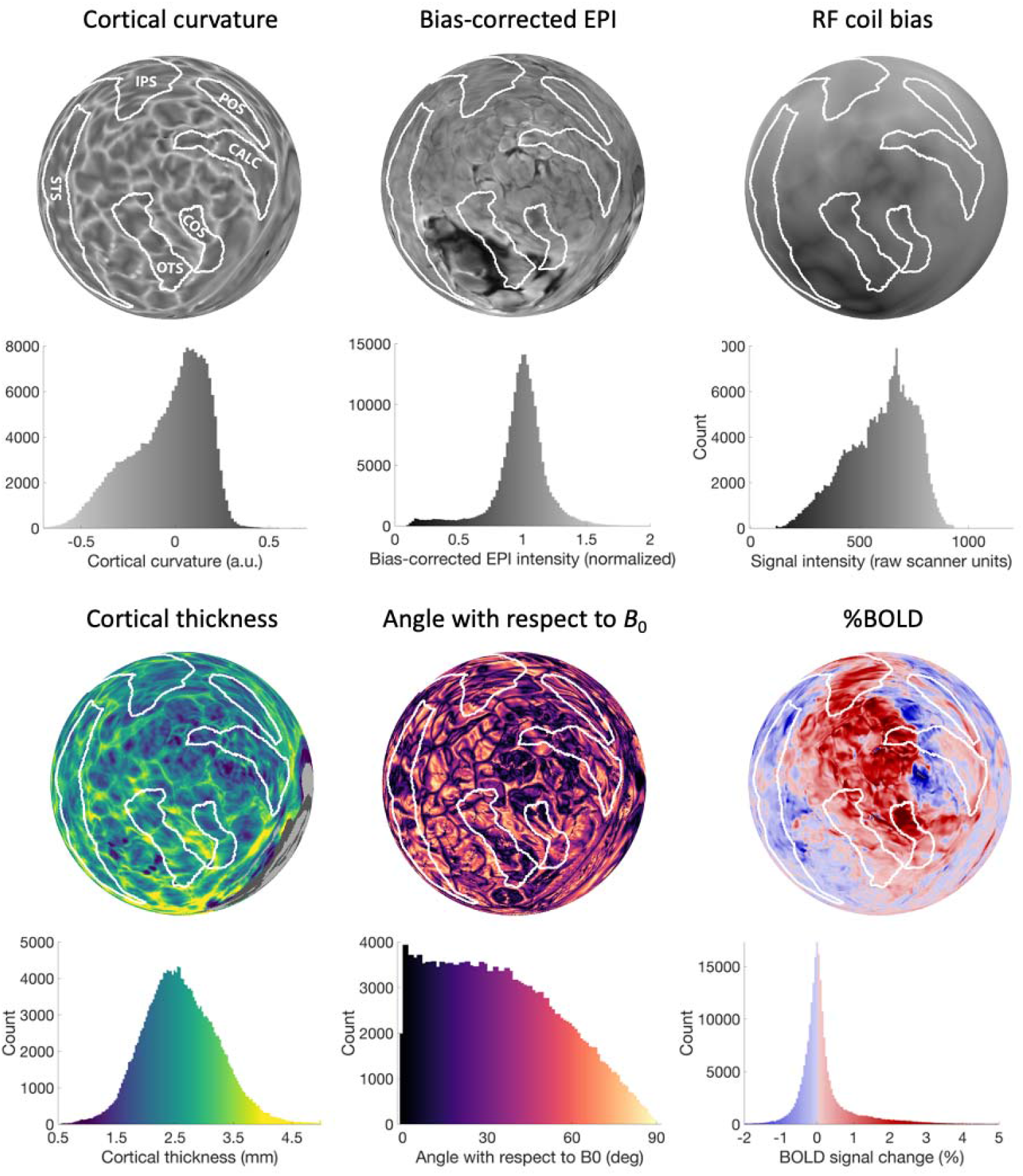
Variation in anatomical and acquisition factors across cortex. Each sphere shows data mapped on the left hemisphere for subject S1 in the NSD dataset. Below each surface map is a histogram of the plotted values. White outlines indicate major cortical sulci. %BOLD represents the average response to the natural scene stimuli used in the NSD dataset. Some of the spatial variability in %BOLD might be due to variability in the depicted non-neural factors.

To understand the potential relationships amongst these five identified factors and %BOLD, we first performed voxel-wise correlation analyses. For these analyses, we used the TDM dataset, as its high spatial resolution facilitates the identification of vascular effects (Kay et al., 2019). We examined data from V1–V3 where neural activity magnitudes can be expected to be relatively homogeneous (although biases were reported before; Liu et al., 2006) given the simple contrast patterns used. In **Figure 6A**, we show pairwise correlations across these five quantities, as well as retinotopic polar angle preference (rescaled between 0 and 90; see Methods) and cortical depth. We find that %BOLD correlates substantially with four factors: curvature (*r* = 0.26), thickness (*r* = −0.17), mean bias-corrected EPI intensity (*r* = −0.25), and depth (*r* = −0.27). We do not find a strong correlation between %BOLD and polar angle. Although results from **Figure 2C–E, Figure 3C** and **Figure 4** suggest a strong negative correlation, the previous analysis included data only from V1. Here we analyze vertices from V1-V3 where this relationship becomes weaker (*r* = −0.05). Overall, we can summarize as follows: %BOLD extracted from V1–V3 tends to be higher at locations that correspond to sulci, in thinner parts of the cortex, in voxels with lower mean bias-corrected EPI intensities, and at more superficial depths.

**Figure 6.**
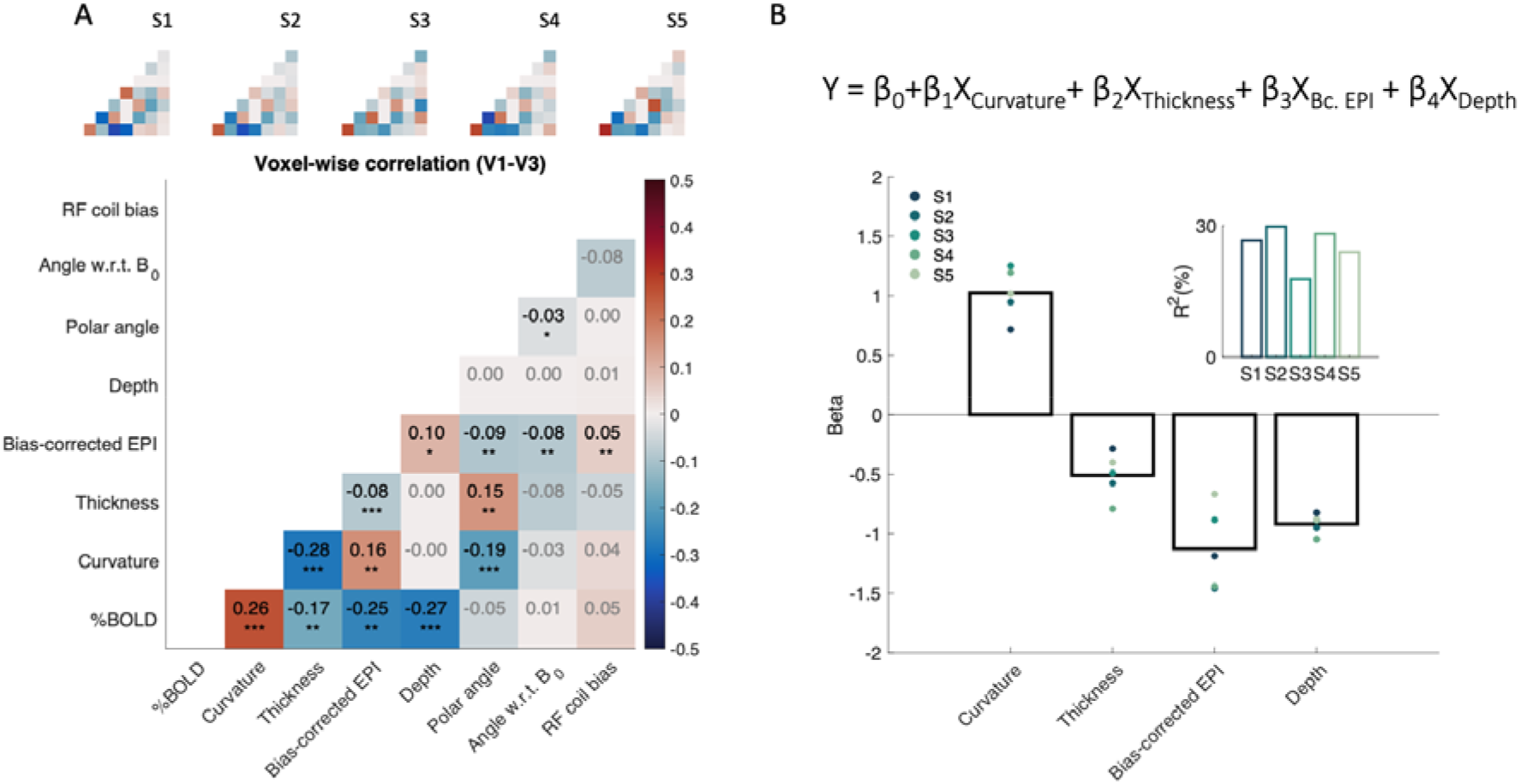
Modeling variations in BOLD signal magnitude. **A)** Correlation (Pearson’s *r*) between a variety of factors and %BOLD extracted from V1–V3 from the TDM dataset. Main plot shows results from data concatenated across all subjects, while inset plots show results from individual-subject data. P-values indicate significance of one sample t-test across subjects; *\*p* < 0.05; *\*\*p* < 0.01; *\*\*\*p* < 0.001. **B)** Regression model for %BOLD. Based on the results of panel A, we selected curvature, thickness, depth and mean bias-corrected EPI as the main non-neural factors that confound %BOLD. These four factors were then used in a multiple linear regression model to predict %BOLD (top). The amount of variance explained by the model is shown in the inset.

Examination of correlations amongst factors yields additional insights **(Figure 6A)**. The strongest correlation that we find is between curvature and thickness (*r* = −0.28), indicating that sulci tend to be thin. Curvature is correlated with mean bias-corrected EPI (*r* = 0.16) and with polar angle (*r* = −0.19), and thickness is correlated with polar angle (*r* = 0.15). Our interpretation of these effects is that venous effects tend be stronger in gyri (consistent with previous findings in Kay et al., 2019), and that the correlations related to polar angle simply reflect the tendency for horizontal meridian representations to fall on sulci (e.g. the calcarine sulcus). Overall, these complex relationships suggest that making sense of non-neural influences on %BOLD requires a broad perspective that considers multiple factors.

### Correcting BOLD signal magnitude for non-neural factors

We now explore whether we can develop a statistical model to compensate for the influence of non-neural factors on %BOLD. We operate under the assumption that any observed correlation between the factors and %BOLD is incidental and does not reflect genuine neural activity variation. Our model is a multiple regression model **(Figure 6B, top)** that uses the main factors of curvature, thickness, depth and mean bias-corrected EPI intensity as continuous variables and attempts to determine a weighted sum of these factors that optimally accounts for variations in %BOLD across cortical locations (see Methods for details).

Fitting the model, we find a strong positive contribution of curvature and negative contributions of thickness, mean bias-corrected EPI intensity and depth **(Figure 6B, bottom)**, consistent with the earlier voxel-wise correlation analyses. Estimated beta weights are fairly consistent across subjects, and the model on average across subjects explains 26% of the variance in %BOLD. A multiple regression model using all 6 factors (adding RF coil bias and angle with respect to *B*_0_) resulted in only minimally larger explained variance, 27% vs. 26%, consistent with the earlier correlation analyses indicating that RF coil bias and angle with respect to *B*_0_ bear little or no relationship with %BOLD.

To better understand the relationship between the identified non-neural factors and %BOLD, we construct a 2D histogram relating the model fit (BOLD prediction based on non-neural factors obtained by multiplying the design matrix and estimated beta weights) and %BOLD **(Figure 7A)**. This reveals a clear nonlinear relationship. To accommodate this nonlinearity, we fit a nonlinear function relating the linear model fit and %BOLD (blue line in **Figure 7A)**. Finally, we remove the contribution of non-neural factors by dividing %BOLD observed at each cortical location by the fit of the nonlinear model. We divide %BOLD by the model fit rather than subtracting the model fit, as we believe that the influence of non-neural factors on %BOLD might impose a type of ‘gain’ field on fMRI responses observed in a given experiment. For example, if there is an excess of macrovasculature in a voxel, we would expect the overall amplitude of the BOLD response from the voxel to be scaled. Note that our method of rescaling BOLD magnitudes does not change the pattern of responses across different experimental conditions within a voxel (while a subtractive approach would). For example, if the response to condition A is 25% higher than the response to condition B, this will continue to be the case after rescaling.

**Figure 7.**
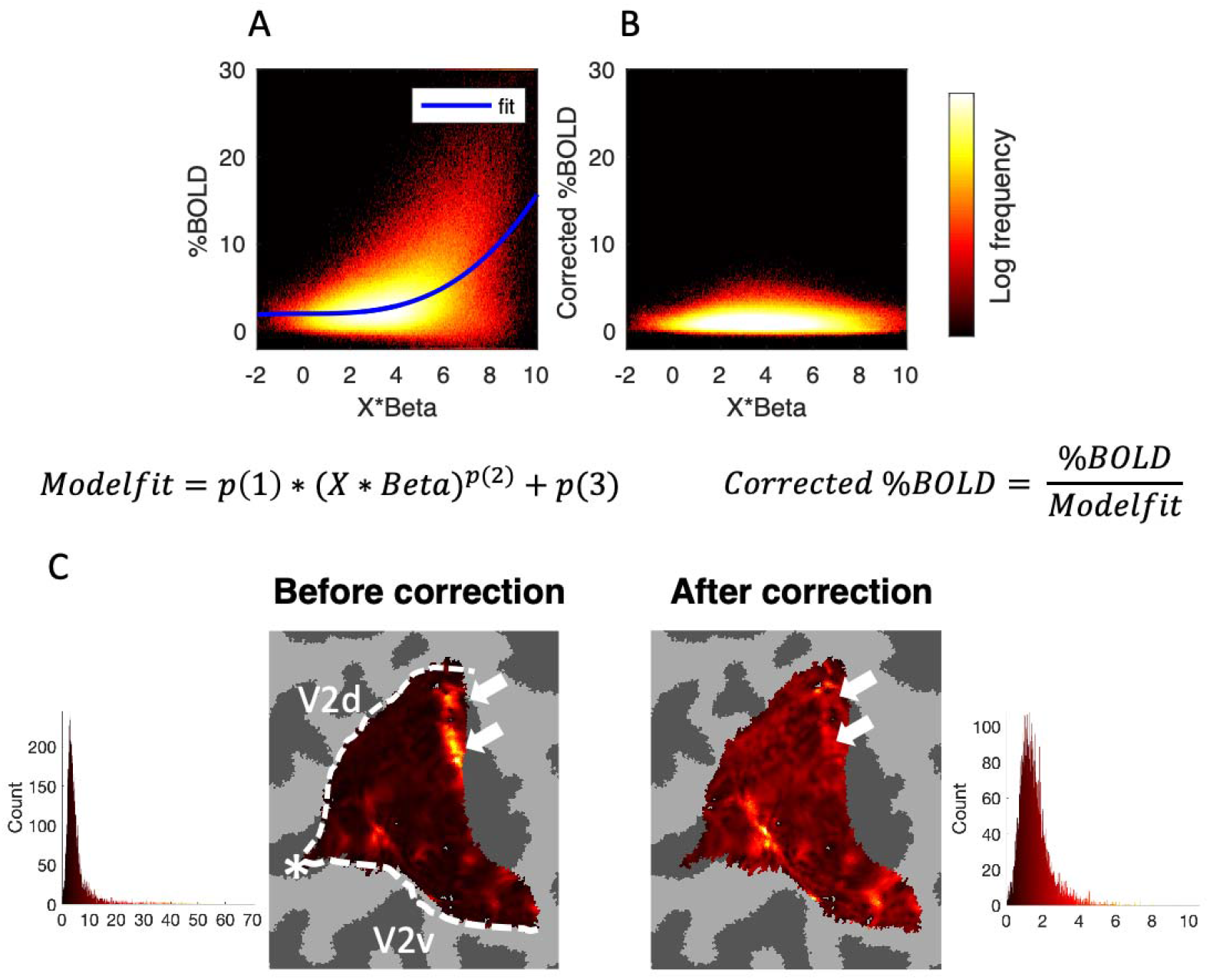
Correction of V1 BOLD inhomogeneity. **A)** Removal of non-neural factors. First, linear combinations of non-neural factors are used to predict %BOLD within V1–V3 using the TDM dataset. The model is fit on data concatenated from all 5 TDM subjects. The model is augmented with a nonlinear power-law function (blue line), which is controlled by a gain parameter (*p*(1)), an exponent parameter (*p*(2)), and a constant term (*p*(3)). **B)** Each voxel’s BOLD responses are divided by the model fit, yielding the corrected %BOLD. **C)** BOLD signal magnitude within V1 before and after the correction (TDM dataset, subject S3, most superficial depth). Asterisk indicates the fovea and dashed lines indicate the boundary between V1 and V2. After correction, some vertices with very high BOLD are eliminated (see white arrows). Within each plot, the color range extends from 0 to the maximum. Each map has an associated histogram that shows all values extracted from V1.

The result of the proposed correction procedure is shown in **Figure 7B**. We see that after the correction procedure, the distribution of BOLD response becomes flatter, indicating the efficacy of the procedure. (Note that what is important is the shape of the distribution of the values, not necessarily the magnitudes of the values.) Increased homogeneity of BOLD magnitude is also visible on the cortical surface **(Figure 7C)**.

To understand whether our method generalizes across datasets, we used the same procedure and performed correction on the NSD dataset. We summarize the effect of the correction by showing the correlations between %BOLD and non-neural factors before and after the correction **(Figure 8A)**. The pattern of results before correction **(Figure 8A, top)** is consistent across the TDM and NSD datasets, except for the reduced correlation with bias-corrected EPI in NSD (see Discussion). Importantly, correlations after the correction are substantially reduced, indicating the efficacy of the method.

**Figure 8.**
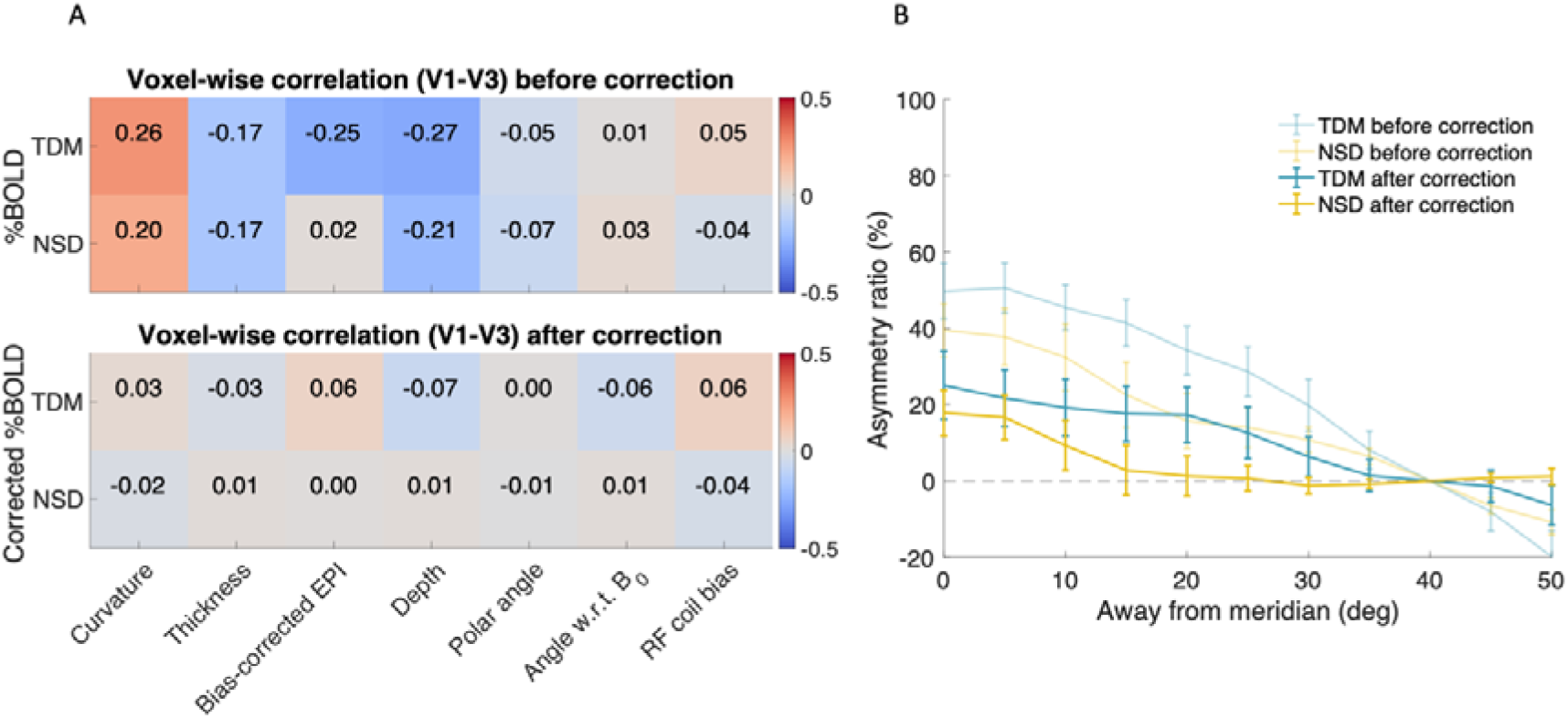
The effect of BOLD inhomogeneity correction. **A)** Voxel-wise correlation between the various factors and %BOLD before and after correction. After correction, correlations are reduced, indicating that the corrected data are less influenced by the non-neural factors. **B)** Dependence of %BOLD on polar angle in V1 before and after the correction for TDM dataset and NSD datasets. The asymmetry drops by about half.

To check whether accounting for non-neural factors increases the homogeneity of BOLD, we quantified the variation of BOLD magnitudes across V1 before and after the correction. Variation was quantified using the semi-interquartile range divided by the median (SIR). Intuitively, if the spread of BOLD magnitudes is small (i.e., %BOLD is relatively homogeneous), SIR will be low, whereas if the spread of BOLD magnitudes is large (i.e., %BOLD is relatively homogeneous), SIR will be high. We find that across subjects, the SIR decreases from 0.42 before correction to 0.34 after correction for TDM and decreases from 0.48 to 0.42 for NSD.

We now return to the experimental effect that motivated this study, namely, BOLD response asymmetries across the horizontal and vertical meridians in V1. We quantify the asymmetry before and after correction **(Figure 8B)**. We find that after accounting for the non-neural factors, the asymmetry drops for TDM from 49.2% to 25.1% and from 40% to 18% for NSD. We thus suggest that some of the observed differences in BOLD response magnitudes are due to non-neural factors.

In the results demonstrated in this paper, the correction method reduces inhomogeneities between the horizontal and vertical meridians. But more generally, it is possible that in other datasets, the method may reveal activity differences that are masked by non-neural factors. For example, voxel A might have a lower neural response than voxel B, but voxel A might reside close to a large vein which would tend to increase %BOLD. In conventional fMRI analyses, both voxels might show similar BOLD magnitude, even though the underlying neural activity is different. The methods proposed in this paper can be viewed as an attempt to obtain better estimates of underlying neural activity.

## Discussion

In this paper, we used three publicly available datasets to assess the degree of homogeneity of BOLD signal magnitude in primary visual cortex. We found that stimulus-evoked BOLD responses, expressed as percent signal change, are up to 50% stronger along the horizontal meridian than the vertical meridian. To investigate whether these magnitude differences can be attributed to differences in local neural activity, we systematically evaluated the potential contribution of several non-neural factors to the observed effect. We found that BOLD signal magnitude correlates with curvature, thickness, depth and macrovasculature (as indexed by bias-corrected EPI intensities). Using a regression-based correction procedure, we were able to increase the homogeneity of BOLD signal magnitude and found that the meridian differences were reduced by half.

### Spatial variations in BOLD magnitude

This study tackles the issue of the neural basis of variation in BOLD signal magnitude. Specifically, we address variation in BOLD across cortical locations for a fixed experimental manipulation, as opposed to variation in BOLD across experimental manipulations for a fixed cortical location. The latter has been heavily studied (Heeger et al., 2000; Logothetis et al., 2001; Heeger and Ress, 2002; Logothetis and Wandell, 2004; Mishra et al., 2021), whereas the former has not yet been systematically studied to the best of our knowledge. If there are indeed non-neural factors that influence BOLD signal variation, taking this into account is critical when interpreting differences in fMRI responses across brain regions.

We acknowledge that a challenge in understanding the neural basis of the BOLD signal is that directly comparable ground-truth measurements of neural activity are typically not available. Moreover, the BOLD signal only indirectly measures the neural response, and its magnitude likely depends on many aspects of neural activity. Increased BOLD signal might be a consequence of more neurons firing, more spikes per neuron, changes in neural correlation, changes in subthreshold activity, and/or changes in what kinds of neurons are most active. Our approach currently does not try to distinguish amongst these causes.

In our analyses, we relied on the working assumption that the experimental paradigms of the three datasets (combined with suitable averaging and analysis procedures) are expected to generate relatively homogeneous patterns of neural activity in early visual cortex. Of course, this may not be exactly the case.

### Non-neural factors that affect BOLD magnitude

#### Mean bias-corrected EPI

Mean bias-corrected EPI is a convenient marker for macrovascular contributions to the fMRI signal (Kay et al., 2019). Vertices contaminated by venous effects show lower intensity values in mean EPI images and often result in high %BOLD magnitude.

In the TDM dataset, we found this to be the case and were able to remove, to some extent, venous effects using the described correction method. We did not, however, find a strong relationship between mean bias-corrected EPI and %BOLD magnitude in the NSD dataset. We suggest that the reason for this apparent discrepancy is that effective discovery of venous contributions requires high-resolution data where voxel size approaches the scale of 1 mm or better. Another important issue to consider is the cerebral sinuses. The sinuses are the largest veins that drain blood from the brain and they exert major effects at certain specific cortical locations. Complicating matters is the fact that the sinuses also produce low EPI intensity, but instead of boosting BOLD magnitude they seem to reduce it, resulting in low %BOLD (Winawer et al., 2010; Jamison et al., 2017). In the present study, we do not attempt to isolate or analyze the effects of the cerebral sinuses, though preliminary analyses indicate that the sinuses do not provide a simple explanation of the horizontal/vertical asymmetry (data not shown).

#### Cortical anatomy

We find that curvature and thickness correlate with BOLD signal magnitude (see **Figure 6A)**. It is known that many anatomical properties vary with thickness and with curvature (Jiang et al., 2021): (i) total neuron count is higher in gyri than it is in sulci (Hilgetag and Barbas, 2005), (ii) gyri tend to be thicker than sulci (Welker, 1990; Hilgetag and Barbas, 2005), (iii) venous effects (resulting in higher BOLD signal amplitude) are more prominent in gyri than they are in sulci (Kay et al., 2019); and (iv) there may even be intrinsic causal relationships between curvature and thickness during anatomical development (Hilgetag and Barbas, 2005). However, the exact anatomical and biophysical mechanisms that might link curvature and thickness to BOLD signal magnitudes are largely unknown, to our knowledge. This is an important issue for future research. Here, we operate under the working assumption that correlations between the BOLD signal and curvature or thickness reflect incidental factors unrelated to local neural activity. We therefore assume that a correction which removes their influence from the BOLD signal is desirable.

#### Orientation of pial veins

It has been reported that regions where the cortical surface is oriented perpendicular to the main magnetic field produce lower BOLD signal than regions where the surface is oriented parallel (Gagnon et al., 2015a; Fracasso et al., 2018). The proposed explanation is that this effect is caused by the orientation of pial veins, which lie parallel to the cortical surface. Our analyses did not replicate this result and indicated little relationship between BOLD magnitude and angle with respect to *B*_0_ (see **Figure 6A)**. One possible explanation could be related to our pre-processing approach, in which fMRI signals are sampled specifically in the gray matter and away from the pial veins that reside on top of the gray matter. This may have dampened effects related to the pial veins. Nonetheless, the prior literature would have predicted some *B*_0_ effect even at inner cortical depths (Viessmann et al., 2019). Alternatively, it is possible that the orientation effects depend in some way on pulse sequence parameters, or the specific brain area being studied. A detailed examination of different datasets would be necessary to resolve these discrepancies.

#### RF coil effects

Due to cortical folding, gyri tend to be closer to the RF coil than sulci. Locations that are further from the coil might have lower mean signal intensities and therefore lower SNR (Srirangarajan et al., 2021), but this should not affect BOLD *magnitudes* expressed in terms of percent signal change. We are not aware of any mechanism that would alter the percent signal change in brain locations that are further away from the RF coil. Indeed, we did not find any relationship between RF coil bias and BOLD magnitude (see **Figure 6A)**.

### Correction for the impact of non-neural factors

Our results show that voxel-wise %BOLD is likely contaminated by several non-neural factors.

To account for these factors, we developed a regression-based correction method. The goal of this method was to introduce a simple, data-driven approach that can be applied irrespectively of the specific experiment or brain region that is under consideration. The underlying premise of the method is that by removing the contribution of non-neural factors, the resulting measures would constitute a better representation of the underlying neural activity. After application of the method, we found that %BOLD becomes more homogenous and correlations between %BOLD and non-neural factors become significantly reduced. Thus, our results indicate that some variation in %BOLD that might be interpreted as change in neural activity likely reflects the variation of non-neural factors.

We believe the results presented in this paper constitute a first step towards developing a cogent strategy for compensating for non-neural biases in BOLD signal magnitudes. Suppressing the influence of non-neural factors has potential applications in pre-surgical planning, where fMRI is routinely used to map motor, speech, and visual areas. The value of fMRI for presurgical planning is currently limited by the accuracy of localizing neural responses (Silva et al., 2018a). BOLD-derived maps that are a better representation of neural activity could lead to more accurate neurosurgical interventions.

It remains to be seen whether the remaining asymmetry across the horizontal and vertical meridians in V1 is a result of genuine neural activity differences, or an effect of other non-neural factors that we were unable to quantify in the present study (which might require additional MRI acquisition measures and/or higher resolution data). It is conceivable that genuine neural activity differences may exist across the horizontal and vertical meridian locations in V1. For example, there is greater cortical magnification along the horizontal than vertical meridian (Silva et al., 2018b; Benson et al., 2021; Himmelberg et al., 2021; Himmelberg et al., 2022), and it is plausible that this might be accompanied by differences in the strength of neural responses.

Although our method is aimed towards more meaningful quantification of the BOLD signal, it differs conceptually from quantitative BOLD (qBOLD) approaches (He and Yablonskiy, 2007; Yablonskiy et al., 2013; Cherukara et al., 2019). On the one hand, qBOLD attempts to model the BOLD signal in terms of its underlying metabolic and hemodynamic components (e.g., blood flow, blood volume, oxygenation extraction), and this in principle may yield measures more closely related to neural activity. On the other hand, the approach we have taken in this paper is to apply analytic methods to BOLD data that consider inhomogeneities that may exist across the brain, with the goal of better estimating local neural activity. Note that the two approaches are not mutually exclusive: one might imagine assessing whether the magnitude of qBOLD measures co-vary with non-neural factors across the brain.

There are other methods that can be used to suppress the contribution of non-neural factors to BOLD signal magnitudes. By identifying early and late components of evoked hemodynamic responses, a temporal decomposition method can be used to estimate BOLD response components more closely linked to the microvasculature, which presumably more closely reflect local neural activity (Kay et al., 2020). Another analysis method focuses on BOLD fluctuations where estimates of slow oscillations (< 0.1 Hz) are used to suppress vascular-related effects (Kazan et al., 2016). Similarly, some methods use the amplitude of fluctuations in resting-state data to rescale the BOLD signal (Di et al., 2013; Guidi et al., 2020). Finally, acquisition methods, such as spin-echo pulse sequences, can be used to suppress unwanted venous effects. Note that all these methods concern effects of the macrovasculature, but systematic biases in BOLD signal magnitudes may in theory persist even if BOLD responses were fully restricted to the microvasculature. Further investigation is necessary to resolve these possibilities.

## Acknowledgments

This research was funded by the US National Eye Institute R01-EY027964, R01-MH111417 and R01-EY027401 to JW. Collection of the NSD dataset was supported by NSF CRCNS grants IIS-1822683 (to KK) and NSF IIS-1822929.

## Author Contributions

J.W.K, K.K. and J.W. conceived and designed this study. J.W.K and K.J. performed data analyses, with K.K. and J.W. providing guidance. J.W.K., J.W, O.F.G and K.K. wrote and edited the manuscript.

